# The Genetic Basis of Incipient Sexual Isolation in *Drosophila melanogaster*

**DOI:** 10.1101/2024.03.01.582979

**Authors:** Akihiko Yamamoto, Wen Huang, Mary Anna Carbone, Robert R. H. Anholt, Trudy F. C. Mackay

## Abstract

Speciation is a fundamental evolutionary process, but the genetic changes accompanying speciation are difficult to determine since true species do not produce viable and fertile offspring. Populations of the same species that are that are partially reproductively isolated are incipient species that can be used to assess genetic changes that occur prior to speciation. *Drosophila melanogaster* from Zimbabwe, Africa are genetically differentiated and partially sexually isolated from cosmopolitan populations worldwide: cosmopolitan males have poor mating success with Zimbabwe females. We used the cosmopolitan *D. melanogaster* Genetic Reference Panel (DGRP) to show there is significant genetic variation in mating success of DGRP males with Zimbabwe females, map genetic variants and genes associated with variation in mating success and determine whether mating success to Zimbabwe females is associated with other quantitative traits previously measured in the DGRP. We performed three genome wide association analyses: for the DGRP lines, for selected flies with high or low mating success from an advanced intercross population (AIP) derived from DGRP lines, and for lines derived from 18 generations of divergent selection from the AIP for mating success with Zimbabwe females. The basis of incipient sexual isolation is highly polygenic and associated with the common African inversion *In(3R)K* and the amount of the sex pheromone 5,9-heptacosadiene in DGRP females. We functionally validated the effect of eight candidate genes using RNA interference. These candidate gene and variant associations provide testable hypotheses for future studies investigating the molecular genetic basis of incipient sexual isolation in *D. melanogaster*.

## Introduction

The genetic basis of speciation – the genetic changes causing the splitting of a panmictic population into two reproductively isolated species – is difficult to determine, since, by definition, complete reproductive isolation is refractory to genetic mapping. Many insights have been gained by mapping the genetic basis of divergence between closely related species with incomplete reproductive isolation, at least in laboratory settings (Coyne and Orr 2004). However, in most cases the mapping resolution is at the level of chromosomes or chromosome regions, and the genetic differentiation includes changes that occurred following speciation as well as those causing or accompanying speciation.

The discoveries that populations of *Drosophila melanogaster* from Zimbabwe (Z), Africa are genetically differentiated from cosmopolitan (C) populations worldwide (Begun and Aquadro 1993) and that there is partial sexual isolation between Z and C populations (Wu et al. 1995; Hollocher et al. 1997a; 1997b) present an ideal scenario to investigate the early stages of sexual isolation – thought to be the first stage in speciation (Ritchie 2007) – in a genetically tractable model system. The partial sexual isolation between Z and C populations is asymmetric and is driven by female choice: Z females do not mate with C males, but all other combinations of mating pairs are successful (Wu et al. 1995; Hollocher et al. 1997a; 1997b). Chromosome substitution analyses indicated that genes contributing to the asymmetric sexual isolation were autosomal, with a larger contribution from the third than the second chromosome (Wu et al. 1995; Hollocher et al. 1997a; 1997b). Mapping factors on the third chromosome by linkage to visible markers identified three regions associated with Z female mating preference (Ting et al. 2001). However, further high-resolution mapping was stymied by the large number of segregating inversions between Z and C populations (Aulard et al. 2002).

Two strategies have been used to gain insight into the genetic differences associated with the partial sexual isolation between Z and C populations. One is association mapping using variants in candidate genes that are divergent between the two populations, and the other is searching for traits that are genetically correlated with the difference in mating behavior between Z and C populations. these strategies were initially applied to differences in cuticular hydrocarbon (CHCs) profiles between the two populations (Fang et al. 2002; Greenberg et al. 2003; Grillet et al. 2012), since CHCs are used as mating cues in both sexes and are divergent between Z and C populations. However, these studies were limited by using small numbers of Z and C strains.

Here, we extend the genetic and trait association designs to the 205 inbred, sequenced C lines of the *D. melanogaster* Genetic Reference Population (DGRP) (Mackay et al. 2012; Huang et al. 2014) and an outbred advanced intercross population (AIP) derived from a subset of DGRP lines. We found significant genetic variation for the mating behavior of Z females with the DGRP (C) males and performed genome-wide association (GWA) mapping of variants associated with DGRP male mating ability with Z females. Since the DGRP has been assessed for over 100 quantitative traits, including CHC composition and ecologically relevant traits (Mackay and Huang 2018), we were also able to assess correlations of male DGRP traits with Z female mating behavior. We used the AIP population to select for C males with increased mating to Z females and performed whole-genome sequencing to identify alleles associated with increased Z female preference. We then used RNA interference (RNAi) to functionally assess the genetic associations at the level of candidate genes. We found that the genetic basis of incipient sexual isolation between DGRP males and Z30 females is highly polygenic and associated with the presence of the common African polymorphic inversion *In(3R)K* as well as the amount of the CHC 5,9-heptacosadiene DGRP females. We identified many candidate genes and variants in the DGRP associated with mating behavior with Z30 females that can be used in future studies of the molecular genetic basis of incipient sexual isolation in *D. melanogaster*.

## Materials and Methods

### Drosophila stocks

The 205 inbred, sequenced DGRP lines were derived from inseminated females collected in Raleigh, NC USA (Mackay et al. 2012; Huang et al. 2014). The Z30 strain from Zimbabwe was a gift from Dr. C. F. Aquadro, Cornell University. Oregon and Samarkand are common C wild type stocks, unrelated to the DGRP lines. RNAi lines were purchased from the Vienna *Drosophila* Resource Center. The *GAL4* driver lines *Act-GAL4* (*P{Act5C-GAL4}25FO1*) and *Ubi-GAL4* (*P{Ubi-GAL4}2*) were obtained from the Bloomington *Drosophila* Stock Center and their major chromosomes that do not contain the drivers were replaced with *Canton-S-B* chromosomes (*CSB, w*^1118^, Yamamoto *et al*. 2008) to minimize background genotype effects. A new driver stock, *Ubi-GAL4[156]*, was created by introducing the original *Ubi-GAL4* transgene onto the third chromosome of *CSB* by *Δ2-3* transposase-mediated hopping.

The AIP was constructed from 40 DGRP lines: DGRP_208, DGRP_301, DGRP_303, DGRP_304, DGRP_306, DGRP_307, DGRP_313, DGRP_315, DGRP_324, DGRP_335, DGRP_357, DGRP_358, DGRP_360, DGRP_362, DGRP_365, DGRP_375, DGRP_379, DGRP_380, DGRP_391, DGRP_399, DGRP_427, DGRP_437, DGRP_486, DGRP_514, DGRP_517, DGRP_555, DGRP_639, DGRP_705, DGRP_707, DGRP_712, DGRP_714, DGRP_730, DGRP_732, DGRP_765, DGRP_774, DGRP_786, DGRP_799, DGRP_820, DGRP_852, DGRP_859. These lines were crossed in a round-robin mating design in generation 1 (*i.e.*, line 1 females by line 2 males, line 2 females by line 3 males, …, line 40 females by line 1 males) to create 40 F1 genotypes. In the second generation, we performed another round-robin cross between pairs of F1 genotypes (*i.e.*, line 1/line 2 F1 females by line 3/line 4 F1 males) to create a highly heterozygous population. At generation 3, 10 replicate populations were established, each with one female and one male from each of the 40 Generation 2 crosses, and flies were allowed to lay eggs for 2 days to minimize natural selection via larval competition. The AIP was maintained from generation 4 in 10 bottles with four females and four males from each of the 10 bottles of the previous generation, for a census population size of 800. All stocks were raised on standard cornmeal/molasses/agar medium at 25°C.

### Mating assay

Five virgin Z30 females and 10 C males were placed without anesthesia in a vial (25 mm diameter x 95 mm high) with 1 ml of medium. All flies were between 4-7 days old. Mating was observed directly and the time to copulation recorded. Each copulating couple was immediately removed using a mouth aspirator through a slit window in a sponge plug (Kitagawa 1979). Mating was observed for 30 minutes, one hour and/or two hours, depending on the experiment. All assays were conducted between 8am to 11am under full lighting at 25°C.

### Quantitative genetics of male mating success with Z30 females in the DGRP

We partitioned the phenotypic variance in male mating success with Z30 females by a mixed model in which the response variable was the mating success rate and the independent variable was the genotype of the flies as a random effect. Broad sense heritability (*H^2^*) is estimated as 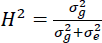, where 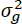 is the variance component due to genotype and 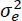 is the variance component due to random environmental effect. Correlations between male mating success and other quantitative traits were computed as Pearson’s correlation coefficient of line means between two traits.

### Genome-wide association (GWA) analysis in the DGRP

We performed a GWA analysis for mating success of DGRP males with Z30 females using the 2,525,695 SNPs and indels with minor allele frequencies greater than 0.02 (Huang et al. 2014). This analysis accounts for effects of Wolbachia infection, cryptic relatedness due to major inversions, and residual polygenic relatedness. In addition, we performed a GWA analysis without correcting for effects of inversions to identify variants within inversions that may contribute to phenotypic variation.

### xQTL mapping

We performed xQTL mapping following a similar procedure as previously described (Huang et al. 2012). A total of 500 male flies at generation 156 were assessed for their mating with Z30 females and 50 fastest maters were selected to form a high mating pool and 50 randomly selected flies were selected to form a control pool. Four biological replicates were performed for each of the high mating and control pools. The pooled flies were sequenced and the sequences were analyzed following a previously described approach (https://github.com/qgg-lab/xqtl). Briefly, reads were mapped to the reference genome and counts of alleles were computed to compare allele frequencies of the high mating and control pool. We used a Z score test in the form of 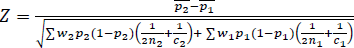 where 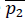 and 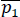 are average allele frequencies in the two groups (high mating and control) across replicates, *n_1_* and *n_2_* are numbers of flies in the pools, and *c_1_* and *c_2_* are the sequencing coverage in the pools. The variance of the allele frequency was averaged with weights (*w_2_*, *w_1_*) according to sequencing depths. *P*-values were obtained by comparing the Z score to standard normal distribution.

### Selection experiments

At AIP generation 30, two replicates of 300 males each were assessed for mating during one hour with Z30 females. Groups of 10 AIP males and five Z30 females were placed in vials, and the first 40 males to mate with Z30 females in each replicate were selected and mated with 40 virgin AIP siblings. Selection was continued for 18 generations. Initially, a single control population was established by observing the mating of 300 AIP males with Z30 females, and randomly selecting 40 males to be parents of the next generation by crossing with 40 AIP females. A second control population was established from the first at generation 5; both were then maintained by scoring mating of 150 males and randomly selecting 40 males to cross with 40 AIP females. At selection Generation 18, the copulation latency of males from the selection and control lines was assessed using F1 hybrid females from a cross between the C strains Oregon and Samarkand. Also at selection generation 18, we paired 600 males from each selection line with 300 Z30 females (60 vials) and collected the first 100 males to mate from each selection line and froze them at -80°C for subsequent DNA sequencing. We also collected and froze 100 randomly selected males from the two generation 18 control populations for subsequent DNA sequencing. The comparison of allele frequencies was done as described above for the xQTL mapping except that the control population and the selected population were not paired, we therefore performed comparisons for all possible pairs and stringently required that the minimal difference was above the threshold.

### DNA sequencing

We sequenced one sample each containing pools of 100 males from the two replicate control and selection lines at selection generation 18 and four samples each containing pools of 50 males from each of the high and control single generation selection experiments. We homogenized the flies from each sample in Gentra Puregene Cell Lysis Solution (Qiagen) with ceramic beads using the TissueLyser (Qiagen Inc.). Genomic DNA was extracted using the Gentra Puregene Tissue Kit (Qiagen) and further purified with AMPure XP magnetic beads (Beckman Coulter). Genomic DNA was fragmented to 300-400bp using ultrasonication (Covaris S220). Fragmented DNA was used to produce barcoded DNA libraries using NEXTflex™ DNA Barcodes (Bioo Scientific, Inc.) with either the TruSeq® DNA Library Prep Kit (Illumina, Inc.) (selection Generation 18 samples) or an Illumina TruSeq compatible protocol (single generation selection samples). Libraries were quantified using Qubit dsDNA HS Kits (Life Technologies, Inc.) and Bioanalyzer (Agilent Technologies, Inc.) to calculate molarity. Libraries were then diluted to equal molarity and re-quantified. The four selection Generation 18 samples were pooled together; and the eight single generation selection samples were pooled together. Pooled library samples were quantified again to calculate final molarity and then denatured and diluted to 14pM. Pooled library samples were clustered on an Illumina cBot. The selection Generation 18 samples were sequenced on one HiSeq2000lane using 100 bp paired-end v3 chemistry, and the eight single generation selection samples were each sequenced on two Hiseq2500 high throughput lanes using 125 bp paired-end v4 chemistry.

### RNA interference (RNAi)

We used three ubiquitously expressed *GAL4* drivers to knock down expression of selected candidate genes: *Act-GAL4*, *Ubi-GAL4*, and *Ubi-GAL4 [156]*. All drivers are in the Canton S B (CSB, a Cosmopolitan strain) genetic background; *Act-GAL4* and *Ubi-GAL4* are maintained over a *CyO* balancer chromosome. We performed luciferase assays to assess the strengths of knock down for each *GAL4* driver. We crossed each driver to a *UAS*-Luciferase stock and collected *GAL4/UAS**-***Luciferase F1 progeny as well as *CyO*/*UAS**-***Luciferase F1 (control) progeny for *Act-GAL4* and *Ubi-GAL4*. The controls for *Ubi-GAL4 [156]* are the CSB/*UAS**-***Luciferase F1 progeny from crossing CSB with the *UAS*-Luciferase stock. We prepared triplicate tissue homogenates from ten F1 progeny from each cross using the Luciferase Cell Culture Lysis 5X Reagent (Promega) to extract total proteins by following the quick-freeze homogenization method outlined by the manufacturer. We quantified the resulting supernatants for their protein concentrations on a SpectraMax M2 (Molecular Devices) using the DC Protein Assay Kit II (BioRad). Luciferase activities were measured on a GloMax Luminometer (Promega) using the Steady-Glo Luciferase Assay System (Promega).

We selected 17 candidate genes to evaluate whether RNAi knockdown of gene expression affected mating performance with Z30 females based on several criteria: low *P*-value of association in any GWA analyses; gene overlap in more than one GWA analysis, functional annotations of candidate genes, and availability of RNAi reagents. We crossed females of each driver line to the RNAi line and the appropriate co-isogenic control line and assessed mating of the F1 males from these crosses with Z30 females at 30 minutes, one hour, and two hours, using 10 replicate vials each with 10 *UAS*-RNAi/*GAL4* males and 5 virgin Z30 females and 20 replicate vials each with 10 control/*GAL4* males and five virgin Z30 females. The exceptions were crosses for *Or67d* RNAi lines, in which the RNAi genotypes were used as male parents and there were 14 replicate vials with 10 males and 5 virgin Z30 females for each of the RNAi and control genotypes. Mating data were analyzed using Fisher Exact tests of mating data for each RNAi line and appropriate control.

## Results

### Variation in DGRP male mating success with Z30 females

We assessed whether the DGRP, a cosmopolitan (C) population, harbored genetic variation in male mating success with Z30 females, which showed a strong preference for Z30 males and avoided mating with C males (Wu et al. 1995; Hollocher et al. 1997a; 1997b). We quantified mating success as the proportion of females that copulated in one or two hours in a no-choice assay in vials with 5 Z30 females and 10 DGRP males. The mean proportion of successful matings within each line varied between 0 and 0.25 after one hour, and 0 and 0.39 after two hours, with respective means of 0.02 and 0.05 (Figure 1, Table S1). There is substantial genetic variation among DGRP males that affect how Z30 females choose their mates. The broad sense heritability (*H*^2^) of “acceptability” of DGRP males to Z30 females was *H*^2^ = 0.27 (*P* = 1.98 x 10^-42^) for the two-hour time point, which was used for all subsequent analyses. The significant genetic variability in acceptability of DGRP males allows us to dissect factors contributing to such variation in the fully sequenced and deeply phenotyped DGRP.

**Figure 1.**
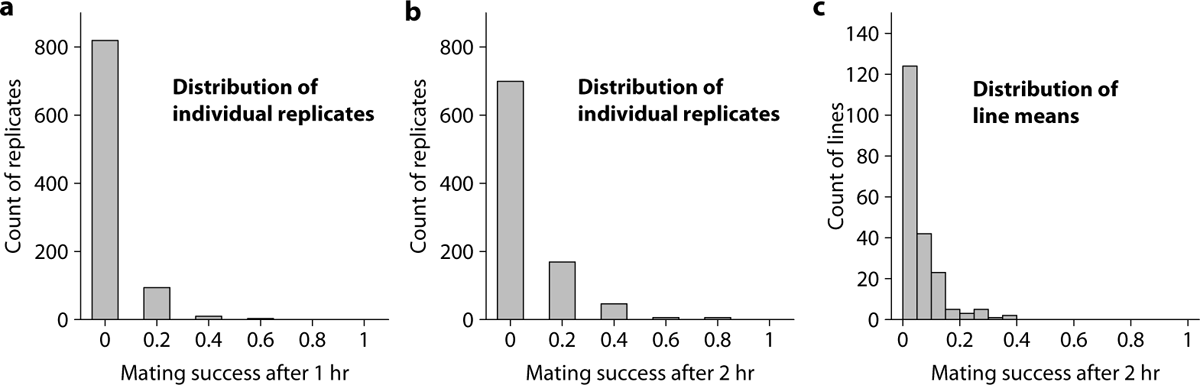
Distribution of DGRP male mating success with Z30 females. (**a**) Distribution of replicate level phenotype for mating success measured after one hour. (**b**) Distribution of replicate level phenotype for mating success measured after two hours. (**c**) Distribution of line means for mating success measured after two hours.

The most abundant CHC moieties in *D. melanogaster* are also sex pheromones and affect mating behavior (Ferveur 1997), and female (Fang et al. 2002; Greenberg et al. 2003) and male (Grillet et al. 2012) CHC composition have been implicated in the mating success of Z30 females. We have previously measured variation in CHC profiles in the DGRP lines (Dembeck et al. 2015) and can thus test these associations in a population with natural variation in CHC abundance. However, we did not find any significant associations of male hydrocarbons, including 7-tricosene and the relative proportion of 7-tricosene (Table S2A), in contrast to a previous report (Grillet et al. 2012). *Desat1* is involved in both the emission and perception of sex pheromones (Bousquet et al. 2012; Nojima et al. 2019). *Desat1* expression is genetically variable in the DGRP, with a broad sense heritability of *H*^2^ = 0.69 in males (Everett et al. 2020). However, variation in male *Desat1* expression is not associated with variation in DGRP male mating success with Z30 females (Table S2B).

Many other quantitative traits that could plausibly be genetically correlated with male mating success with Z30 females are genetically variable in the DGRP and have been measured under the same conditions as this study. We assessed the correlations of male aggressive behavior (Shorter et al. 2015), startle response, starvation resistance and chill coma recovery time (Mackay et al. 2012), phototaxis (Carbone et al. 2016), sleep traits and waking activity (Harbison et al. 2013), DGRP male mating success with C females (Yamamoto et al. 2023), body weight and body size (Everett et al. 2020), food consumption (Garlapow et al. 2015) and metabolic traits (Everett et al. 2020) (Table S2C-K). The only quantitative trait that is significantly, albeit moderately, correlated with DGRP male mating success with Z30 females is DGRP male mating success with Oregon/Samarkand F1 hybrid C females (*r* = 0.256, *P* = 0.00021, Table S2I), suggesting that the male component of the mating success trait is partially independent of females (Yamamoto et al. 2024).

### GWA analyses of male DGRP mating success with Z30 females

*D. melanogaster* populations are polymorphic for many chromosome inversions that often have population specific frequencies between African and C populations (Corbett-Detig and Hartl 2012). Standard and inverted sequences are genetically divergent due to lack of recombination between them (Corbett-Detig and Hartl 2012, Mackay et al. 2012). Therefore, we evaluated whether inversions segregating in the DGRP were associated with DGRP male mating success with Z30 females. We found that *In(3R)K* (proximal and distal breakpoints *3R*_7576289 and *3R*_21966092, respectively), which has a high frequency in African populations but is rare in C populations (Corbett-Detig and Hartl 2012), has a large effect on DGRP mating success with C females (*P* = 9.16×10^-5^, Table S3A, Figure 2a). The *In(3R)K*/ST inversion heterozygotes have the highest proportion of Z30 female matings relative to the standard karyotype (*P* = 1.77×10^-5^). In addition, *In(3R)Mo* (proximal and distal breakpoints *3R*_17232639 and *3R*_24857019, respectively), which is absent in Africa and rare in most C populations (Corbett-Detig and Hartl 2012), but which has a fairly high frequency in the DGRP (Mackay et al. 2012), is also associated with DGRP male mating success with Z30 females (*P* = 0.04, Table S3A, Figure 2b). In this case, it is the homozygous inversion genotype that has the highest proportion of Z30 female matings relative to the standard karyotype (*P* = 1.61×10^-2^). Furthermore, *In(2L)t* (proximal and distal break points *2L*_13154180 and *2L*_2225744, respectively), which is common in African and C populations, is also associated with male mating success (*P* = 0.02, Table S3A, Figure 2c) where the homozygous inversion genotype is associated with higher mating success (*P* = 0.02, Table S3A). These results indicated that the inversions may themselves contain variants that contribute to male mating success. Therefore, we performed two GWA analyses for variants at MAF > 0.02, one accounting for the effect of inversions on the trait (Huang et al. 2014), and one without using inversion status as covariates.

**Figure 2.**
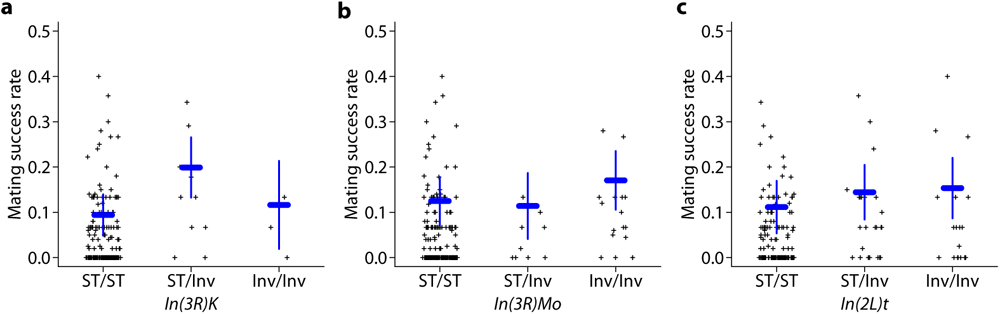
Effects of inversions on DGRP male mating success with Z30 females. (**a**) Effect of *In(3R)K* on mating success. (**b**) Effect of *In(3R)Mo* on mating success. (**c**) Effect of *In(2L)t* on mating success. The blue horizontal bar represents least squares mean of each group and vertical bar represents 95% confidence interval.

The top variants in the GWA analysis (reporting *P*-value < 10^-5^) for which the effects of inversions were accounted for identified 156 variants in or near 115 genes (Tables S3B, S3D). Four intronic variants in three genes (*mew*, *CG40470*, *wry*) were significant after applying a stringent Bonferroni correction for multiple tests (0.05/2,525,695 variants = 1.98 × 10^-8^). The top variants in the GWA analysis for which the effects of inversions were not accounted identified 266 variants in or near 182 genes (Tables S3C, S3D). Six variants in five genes (*mew*, *CG40470*, *CG10226*, *CR44546*, *Or67d*) were significant after applying the Bonferroni correction. A total of 73 genes overlapped between the two GWA analyses, 42 were unique to the analysis corrected for inversions and 109 were unique to the analysis not corrected for inversions (Table S3D). The 224 genes identified in these analyses were enriched (Mi et al. 2020) for Gene Ontology terms involved in cellular signaling, including synaptic signaling and G-protein coupled receptor signaling, and several canonical signaling pathways (Decapentaplegic, Screw, Transforming growth factor beta) (Table S3E).

### Extreme QTL (xQTL) mapping for male mating success with Z30 females

A complementary approach to GWA analysis using the DGRP lines is to use extreme QTL (xQTL) mapping (Ehrenreich et al. 2010) by selecting males from an outbred population that readily mate with Z30 females and comparing their allele frequencies genome-wide with equal numbers of randomly selected males. We constructed a highly heterozygous outbred AIP from a subset of 40 DGRP lines. We sequenced genomic DNA from four pools of 50 males each from the same AIP population at generation 156 that mated rapidly with Z30 females, and four pools of 50 randomly selected males from the same population. We identified 45 variants in or near 44 genes (*P* < 10^-5^) in this analysis (Tables S4A, S4B). Only one gene (*CG42368*) was shared between the xQTL analysis and GWA analysis in the DGRP (Table S4C), which may be due to context dependent genetic effects (Huang et al. 2012).

### Selection from an outbred population for increased Z30 female mating success

Heritable traits are expected to respond to directional artificial selection, during which genetic differentiation is expected to occur. We therefore performed a multi-generation selection experiment and sequenced selected and control populations to identify genomic regions that responded to selection. Unlike xQTL mapping, selection and drift both extend linkage disequilibrium (LD) in selection lines, reducing the resolution of mapping.

We performed 18 generations of selection of C males for mating success to Z30 females from an outbred AIP. The selected lines reached more than 40% mating success in one hour relative to the control lines by generation 18, which had an average mating success of 20% (Figure 3, Table S5). While there was considerable fluctuation, mean deviation of selected lines from contemporary control lines showed a clear directional trend towards higher male mating success with Z30 females. Remarkably, the selection response remained even when mating to a tester C line females. At generation 18, the average proportion of Oregon/Samarkand F1 hybrid females mating with control males was 0.567, while with selected males was 0.729 (Table S5). This result is consistent with the genetic correlation between mating success with Z30 females and that with Oregon/Samarkand F1 females (Table S2I) and also suggesting that the male mating success trait is in part independent of females (Yamamoto et al. 2024).

**Figure 3.**
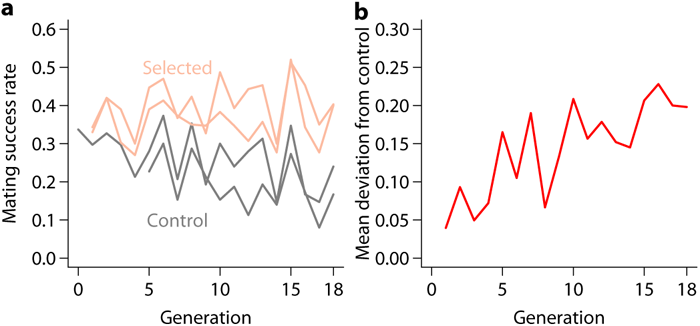
Response to 18 generations of selection from an advanced intercross population derived from 40 DGRP lines for male mating success with Z30 females. (**a**) Phenotypic trends in individual replicate populations. (**b**) Average effect of selection.

To identify genomic regions that responded to selection, we sequenced genomic DNA from pools of 100 males of the replicate control and selected lines. We identified 2,223 variants in or near 968 genes with divergent allele frequencies between the selected and control lines at *P* < 10^-5^ (Tables S6A, S6B). The homozygous DGRP, xQTL mapping in the outbred population and multi-generation selection experiment are complementary and have different strengths and weaknesses. A total of 31 genes were in common between two of these analyses: 23 between the DGRP and multi-generation selection analyses, one between the DGRP and xQTL mapping analyses, and seven between the multi-generation selection and xQTL mapping analyses (Table S4C).

### RNAi of candidate genes

To functionally validate candidate genes identified in these experiments, we utilized RNAi lines to specifically knockdown expression of candidate genes. We first assessed the strength of three ubiquitously expressed *GAL4* drivers (*Act-GAL4*, *Ubi-GAL4*, and *Ubi-GAL4 [156]*) using a luciferase assay. Based on this assay, the relative strengths of the *GAL4* drivers are *Act-GAL4* >*Ubi-GAL4* > *Ubi-GAL4[156]* (Table S7). We chose 17 candidate genes from the DGRP and selection analyses for functional analyses. We included genes that had a small *P*-value from any analysis (*CG33144*, *CG42458*, *CG44837*, *frac*, *mew*, *wry*); were present in two analyses (*btsz*, *CG34114*, *nmo*, *Rbp6*, *tkv*); and had involvement in sensory perception (*dpr1*, *Or67d*), nervous system development and function (*C15*), and/or brain gene expression (*CG1136*, *CG42672*, *jvl*). We evaluated their effects on mating with Z30 females using the three ubiquitously expressed *GAL4* drivers. RNAi of eight of these genes (47.1%) affected mating success of males with Z30 females. RNAi of *CG44837* increased male mating success with Z30 females compared to the control; and RNAi of *btsz*, *C15*, *CG1136*, *CG42672*, *dpr1*, *nmo* and *jvl* decreased male mating success with Z30 females relative to the control (Table S8).

## Discussion

Most previous studies of the genetic basis of the differences in mating behavior of Z and C populations have focused on CHCs that are sex pheromones. The most common female CHC in African and Caribbean populations is 5,9-heptacosadiene, while the most common female CHC in Cosmopolitan populations is 7,11-heptacosadiene. The abundance of 5,9-heptacosadiene has been associated with a regulatory polymorphism in *Desat2*, which has a 16 bp deletion in C populations and an intact allele in Z populations (Fang et al. 2002, Greenberg et al. 2003). Interestingly, there is significant variation in the amount of 5,9-heptacosadiene among DGRP females, with a broad sense heritability of *H*^2^= 0.93 (Dembeck et al. 2015). This might be partially attributable to the presence of the African *Desat2* allele in 17 DGRP lines. However, this allele is not perfectly associated with either variation in the amount of 5,9-heptacosadiene (Dembeck et al. 2015) or mating success with Z30 females (Table S2). For example, DGRP_105 and DGRP_235 both have the African *Desat2* allele; but DGRP_105 has low amounts of 5,9-heptacosdiene and a low proportion of mating with Z30 females; while DGRP_235 has high amounts of 5,9-heptacosdiene and no mating with Z30 females. The correlation between amount of 5,9-heptacosdiene and Z30 mating success in the 17 DGRP lines with the African *Desat2* allele is *r* = 0.077 (*P* = 0.769) (Table S2A). Furthermore, *Desat2* is not expressed in any of the DGRP lines, including those with the African *Desat2* allele (Everett et al. 2021). We performed a GWA analysis of the amount of 5,9-heptacosadiene in DGRP females (Table S9) using the data of Dembeck et al. (2015). The amount of 5,9-heptacosadiene is associated with *In(3R)K* inversion status for both homozygous and heterozygous inversions (Table S9A), and with a total of 612 variants (Table S9B) in or near 341 genes (Table S9C). The African *Desat2* allele was not among the top associated variants. However, 17 genes overlapped between the GWA analyses for Z30 female mating success and amount of 5,9-heptacosadiene in DGRP females (Table S9C), of which three genes (*btsz*, *CG18208* and *Octbeta2R*) are located in *In(3R)K* and are good candidates for the strong association of this inversion with both Z30 mating success and amount of 5,9-heptacosadiene in DGRP females.

We did not find a correlation with Z30 female mating success and the relative amount of 7-tricosene in males, as previously reported (Grillet et al. 2012), nor with expression of *Desat1*, which is involved in both the emission and perception of sex pheromones (Bousquet et al. 2012; Nojima et al. 2019). The failure to replicate these earlier results may be attributable to the larger number of C lines tested in this report. We did find a significant correlation of Z30 female mating success with mating success of DGRP males and Oregon/Samarkand F1 C females (Yamamoto et al. 2022). This correlation may be due to genetic variation in DGRP male mating success in general.

We performed three unbiased GWA analysis screens to detect DGRP genes and variants associated with mating success with Z30 females that have different advantages and disadvantages. The DGRP GWA ANALYSIS is adequately powered to detect common variants with fairly large effects (Mackay and Huang 2018), but not rare variants; and the DGRP has high mapping precision because linkage disequilibrium (LD) declines rapidly with physical distance in this population (Mackay et al. 2012; Huang et al. 2014). The analysis of allele frequency divergence from multiple generations of selection has the power to detect rare variants that increase in frequency in the selection lines, but little power to disentangle the effects of selection and genetic drift for common alleles with only two replicate selection and control lines; further, selection and drift both cause LD, reducing the precision of mapping (Garlapow et al. 2017). The single generation selection experiment can detect common alleles associated with Z30 female mating because the effect of drift and LD is less than the multi-generation selection experiment due to the large effective population size of the AIP, but it has little power to detect rare alleles. Both AIP designs have reduced genetic variation compared to the entire DGRP. Finally, the DGRP lines are inbred while the AIP population is outbred, and there is inbreeding depression for male mating behavior. The mean mating frequency to Z30 females of the DGRP lines used as parents for the AIP is 0.06 (Table S2), while the mean mating frequency to Z30 females of AIP males at generation 0 of the selection experiment is 0.34 (Table S4). Given the contrasting strengths and weaknesses of the three experimental designs, it is not surprising that the overlap between candidate genes identified in each screen is limited. However, it is clear that the genetic architecture of incipient sexual isolation between Z and C *D. melanogaster* strains is highly polygenic.

We functionally assessed the effects of knocking down gene expression of 17 candidate genes using *UAS*-RNAi constructs and three ubiquitous *GAL4* drivers with different strengths. RNAi of one candidate gene (*CG44837*) in C males resulted in increased mating to Z30 females, while RNAi of seven candidate genes (*C15*, *dpr1*, *CG1136*, *CG42672*, *btsz*, *jvl*, *nmo*) in C males resulted in decreased mating to Z30 females. Interestingly, *btsz* and *nmo* are also candidate genes for the amount of 5,9-heptacosadience in C females (Table S9C, Dembeck et al. 2015). *btsz* (*bitesize*) encodes a synaptotagmin-like protein with annotated roles in actin filament organization, apical junction assembly, gastrulation (germ band assembly), morphogenesis of embryonic epithelium, and lumen formation in the tracheal system (Pilot et al. 2006; Jayanandanan et al. 2014). *btsz* has not previously been annotated to affect behavior, but it is expressed in the larval and adult central nervous system (Larkin et al. 2021). *nmo* (*nemo*) encodes a proline-directed serine/threonine kinase with multiple pleiotropic roles in development, including eye (Choi and Benzer 1994) and wing (Verheyen et al. 2001) development and regulation of the Wnt signaling pathway (Verheyen et al. 2001). *nmo* is expressed in adult brains (Larkin et al. 2021) and affects gravitaxis behavior (Toma et al. 2001). *C15* encodes a transcription factor and affects antenna, chaeta, and leg development (Campbell 2005; Kojima et al. 2005). *dpr1* (*defective proboscis extension response 1*) is involved in salt aversion, sensory perception of salty taste (Nakamura et al. 2002) and synapse organization (Carrillo et al. 2015) and is expressed in adult brains (Larkin et al. 2021). *jvl* (*javelin-like*) encodes a microtubule-associated protein which regulates mRNA localization during development, affects chaetae development (Durban-Bar et al. 2011), and is expressed ubiquitously, including the larval and adult central nervous system (Larkin et al. 2021). *CG1136*, *CG42672* and *CG44837* are computationally predicted genes for which there are no experimentally annotated functions, although all are expressed in the adult brain (Larkin et al. 2021). None of these candidate genes has been previously associated with mating behavior, although they are plausible candidates based on brain gene expression.

Functional assessment of phenotypes using RNAi in *D. melanogaster* is facilitated by large numbers of publicly available *UAS*-RNAi stocks and a wide variety of *GAL4* drivers with different expression patterns (Larkin et al. 2021). However, RNAi cannot mimic the effects of candidate SNPs. For example, the effects of intronic SNPs in *mew* (*multiple edematous wings*) and an insertion/deletion polymorphism 47 bp upstream of the transcription start site of *Or67d* (*Odorant receptor 67d*) had among the largest effects and lowest *P*-values in the DGRP GWA analysis (Table S2). RNAi of *mew* resulted in lethality with the stronger *Ubi*-*GAL4* and *Act*-*GAL4* drivers that gave phenotypic effects for other genes. There was no effect of RNAi of *Or67d* with any driver, but it is a particularly interesting candidate gene because it is the receptor for the male-specific sex pheromone, 11-*cis*-vaccenyl acetate, which inhibits male and promotes female mating behavior (Kurtovic et al. 2007). Rigorous functional assessment of the effects of the insertion/deletion polymorphism in *Or67d* or polymorphisms in other candidate genes affecting the mating behavior of C males with Z females would entail either evaluating these associations using in additional DGRP lines that were not used in this study (Baker et al. 2021) or creating scarless allelic replacements of the polymorphic alleles in a common homozygous genetic background. Our results raise the question of how candidate genes expressed in the nervous system along with those associated with sensory perception are functionally interconnected to mediate mating cues and how the genetic underpinnings of such functional ensembles enable the evolutionary trajectory that leads to incipient sexual isolation. The candidate gene and variant associations presented here provide testable hypotheses for future studies investigating the molecular genetic basis of incipient sexual isolation in *D. melanogaster*.

## Notes

### Competing Interest Statement

The authors have declared no competing interest.

## References

Aulard S, David JR, Lemeunier F. 2002. Chromosomal inversion polymorphism in Afrotropical populations of *Drosophila melanogaster*. Genet Res 79: 49–63.

Baker BM, Carbone MA, Huang W, Anholt RRH, Mackay TFC. 2021. Genetic basis of variation in cocaine and methamphetamine consumption in outbred populations of *Drosophila melanogaster*. Proc Natl Acad Sci USA 118: e2104131118.

Begun DJ, Aquadro CF. 1993. African and North American populations of *Drosophila melanogaster* are very different at the DNA level. Nature 365: 548–550.

Bousquet F, Nojima T, Houot B, Chauvel I, Chaudy S, Dupas S, Yamamoto D, Ferveur JF. 2012. Expression of a desaturase gene, *desat1*, in neural and nonneural tissues separately affects perception and emission of sex pheromones in Drosophila. Proc Natl Acad Sci USA 109: 249–254.

Carbone MA, Yamamoto A, Huang W, Lyman RA, Meadors TB, Yamamoto R, Anholt RR, Mackay TFC. 2016. Genetic architecture of natural variation in visual senescence in Drosophila. Proc Natl Acad Sci USA 113: E6620–E6629.

Campbell G. 2005. Regulation of gene expression in the distal region of the Drosophila leg by the Hox11 homolog, C15. Dev Biol 278: 607–618.

Carrillo RA, Özkan E, Menon KP, Nagarkar-Jaiswal S, Lee PT, Jeon M, Birnbaum ME, Bellen HJ, Garcia KC, Zinn K. 2015. Control of synaptic connectivity by a network of Drosophila IgSF cell surface proteins. Cell 163: 1770–1782.

Choi KW, Benzer S. 1994. Rotation of photoreceptor clusters in the developing Drosophila eye requires the *nemo* gene. Cell 78: 125–136.

Corbett-Detig RB, Hartl DL. 2012. Population genomics of inversion polymorphisms in *Drosophila melanogaster*. PLoS Genet 8: e1003056.

Coyne JA, Orr HA. 2004. *Speciation*. Sinauer Associates, Inc. Sunderland MA.

Dembeck LM, Böröczky K, Huang W, Schal C, Anholt RR, Mackay TFC. 2015. Genetic architecture of natural variation in cuticular hydrocarbon composition in *Drosophila melanogaster*. Elife 4: e09861.

Dubin-Bar D, Bitan A, Bakhrat A, Amsalem S, Abdu U. 2011. Drosophila *javelin-like* encodes a novel microtubule-associated protein and is required for mRNA localization during oogenesis. Development 138: 4661–4671.

Ehrenreich IM, Torabi N, Jia Y, Kent J, Martis S, Shapiro JA, Gresham D, Caudy AA, Kruglyak L. 2010. Dissection of genetically complex traits with extremely large pools of yeast segregants. Nature 464: 1039–1042

Everett LJ, Huang W, Zhou S, Carbone MA, Lyman RF, Arya GH, Geisz MS, Ma J, Morgante F, St Armour G, Turlapati L, Anholt RRH, Mackay TFC. 2020. Gene expression networks in the Drosophila Genetic Reference Panel. Genome Res 30: 485–496.

Fang S, Takahashi A, Wu CI. 2002. A mutation in the promoter of *desaturase 2* is correlated with sexual isolation between Drosophila behavioral races. Genetics 162: 781–784.

Ferveur JF. 1997. The pheromonal role of cuticular hydrocarbons in *Drosophila melanogaster*. Bioessays 19: 353–358.

Garlapow ME, Everett LJ, Zhou S, Gearhart AW, Fay KA, Huang W, Morozova TV, Arya GH, Turlapati L, St Armour G, Hussain YN, McAdams SE, Fochler S, Mackay TFC. 2017. Genetic and genomic response to selection for food consumption in *Drosophila melanogaster*. Behav Genet 47: 227–243.

Garlapow ME, Huang W, Yarboro MT, Peterson KR, Mackay TFC. 2015. Quantitative genetics of food intake in *Drosophila melanogaster*. PLoS One 10: e0138129.

Greenberg AJ, Moran JR, Coyne JA, Wu CI. 2003. Ecological adaptation during incipient speciation revealed by precise gene replacement. Science 302: 1754–1757.

Grillet M, Everaerts C, Houot B, Ritchie MG, Cobb M, Ferveur JF. 2012. Incipient speciation in *Drosophila melanogaster* involves chemical signals. Sci Rep 2: 224.

Harbison ST, McCoy LJ, Mackay TFC. 2013. Genome-wide association study of sleep in *Drosophila melanogaster*. BMC Genomics 14: 281.

Hollocher H, Ting CT, Pollack F, Wu CI. 1997a. Incipient speciation by sexual isolation in *Drosophila melanogaster*: variation in mating preference and correlation between sexes. Evolution 51: 1175–1181.

Hollocher H, Ting CT, Wu ML, Wu CI. 1997b. Incipient speciation by sexual isolation in *Drosophila melanogaster*: extensive genetic divergence without reinforcement. Genetics 147: 1191–201.

Huang W, Massouras A, Inoue Y, Peiffer J, Ràmia M, Tarone AM, Turlapati L, Zichner T, Zhu D, Lyman RF, Magwire MM, Blankenburg K, Carbone MA, Chang K, Ellis LL, Fernandez S, Han Y, Highnam G, Hjelmen CE, Jack JR, Javaid M, Jayaseelan J, Kalra D, Lee S, Lewis L, Munidasa M, Ongeri F, Patel S, Perales L, Perez A, Pu L, Rollmann SM, Ruth R, Saada N, Warner C, Williams A, Wu YQ, Yamamoto A, Zhang Y, Zhu Y, Anholt RR, Korbel JO, Mittelman D, Muzny DM, Gibbs RA, Barbadilla A, Johnston JS, Stone EA, Richards S, Deplancke B, Mackay TFC. 2014. Natural variation in genome architecture among 205 *Drosophila melanogaster* Genetic Reference Panel lines. Genome Res 24: 1193–1208.

Huang W, Richards S, Carbone MA, Zhu D, Anholt RR, Ayroles JF, Duncan L, Jordan KW, Lawrence F, Magwire MM, Warner CB, Blankenburg K, Han Y, Javaid M, Jayaseelan J, Jhangiani SN, Muzny D, Ongeri F, Perales L, Wu YQ, Zhang Y, Zou X, Stone EA, Gibbs RA, Mackay TFC. 2012. Epistasis dominates the genetic architecture of Drosophila quantitative traits. Proc Natl Acad Sci USA 109: 15553–15559.

Jayanandanan N, Mathew R, Leptin M. 2014. Guidance of subcellular tubulogenesis by actin under the control of a synaptotagmin-like protein and Moesin. Nat Commun 5: 3036.

Kitagawa O. 1979. D Moriwaki (ed) Shoujoubae no iden jisshu (in Japanese) P. 58, Baifukan.

Kojima T, Tsuji T, Saigo K. 2005. A concerted action of a paired-type homeobox gene, *aristaless*, and a homolog of Hox11/tlx homeobox gene, *clawless*, is essential for the distal tip development of the Drosophila leg. Dev Biol 279: 434–445.

Kurtovic A, Widmer A, Dickson BJ. 2007. A single class of olfactory neurons mediates behavioural responses to a Drosophila sex pheromone. Nature 446: 542–546.

Larkin A, Marygold SJ, Antonazzo G, Attrill H, dos Santos G, Garapati PV, Goodman JL, Gramates LS, Millburn G, Strelets VB, Tabone CJ, Thurmond J, FlyBase Consortium. 2021. FlyBase: updates to the *Drosophila melanogaster* knowledge base. Nucleic Acids Res 49: D899–D907.

Mackay TFC, Huang W. 2018. Charting the genotype-phenotype map: lessons from the *Drosophila melanogaster* Genetic Reference Panel. Wiley Interdiscip Rev Dev Biol 7: 10.1002/wdev.289.

Mackay TFC, Richards S, Stone EA, Barbadilla A, Ayroles JF, Zhu D, Casillas S, Han Y, Magwire MM, Cridland JM, Richardson MF, Anholt RR, Barrón M, Bess C, Blankenburg KP, Carbone MA, Castellano D, Chaboub L, Duncan L, Harris Z, Javaid M, Jayaseelan JC, Jhangiani SN, Jordan KW, Lara F, Lawrence F, Lee SL, Librado P, Linheiro RS, Lyman RF, Mackey AJ, Munidasa M, Muzny DM, Nazareth L, Newsham I, Perales L, Pu LL, Qu C, Ràmia M, Reid JG, Rollmann SM, Rozas J, Saada N, Turlapati L, Worley KC, Wu YQ, Yamamoto A, Zhu Y, Bergman CM, Thornton KR, Mittelman D, Gibbs RA. 2012. The *Drosophila melanogaster* Genetic Reference Panel. Nature 482: 173–178.

Mi H, Ebert D, Muruganujan A, Mills C, Albou LP, Mushayamaha T, Thomas PD. 2020. PANTHER version 16: a revised family classification, tree-based classification tool, enhancer regions and extensive API Nucl Acids Res 49: D394–D403.

Nakamura M, Baldwin D, Hannaford S, Palka J, Montell C. 2002. Defective proboscis extension response (DPR), a member of the Ig superfamily required for the gustatory response to salt. J Neurosci 22: 3463–3472.

Nojima T, Chauvel I, Houot B, Bousquet F, Farine JP, Everaerts C, Yamamoto D, Ferveur JF. 2019. The *desaturase1* gene affects reproduction before, during and after copulation in *Drosophila melanogaster*. J Neurogenet 33: 96–115.

Pilot F, Philippe JM, Lemmers, C, Lecuit T. 2006. Spatial control of actin organization at adherens junctions by a synaptotagmin-like protein Btsz. Nature 442: 580–584.

Ritchie MG. 2007. Sexual selection and speciation. Annu Rev Ecol Evol Syst 38: 79–102.

Shorter J, Couch C, Huang W, Carbone MA, Peiffer J, Anholt RR, Mackay TFC. 2015. Genetic architecture of natural variation in *Drosophila melanogaster* aggressive behavior. Proc Natl Acad Sci USA 112: E3555–E3563.

Ting CT, Takahashi A, Wu CI. 2001. Incipient speciation by sexual isolation in Drosophila: concurrent evolution at multiple loci. Proc Natl Acad Sci USA 98: 6709–6713.

Toma DP, White KP, Hirsch J, Greenspan RJ. 2002. Identification of genes involved in *Drosophila melanogaster* geotaxis, a complex behavioral trait. Nat Genet 31: 349–353.

Verheyen EM, Mirkovic I, MacLean SJ, Langmann C, Andrews BC, MacKinnon C. 2001. The tissue polarity gene *nemo* carries out multiple roles in patterning during Drosophila development. Mech Dev 101: 119–132.

Wu CI, Hollocher H, Begun DJ, Aquadro CF, Xu Y, Wu ML. 1995. Sexual isolation in *Drosophila melanogaster*: a possible case of incipient speciation. Proc Natl Acad Sci USA 92: 2519–2523.

Yamamoto A, Zwarts L, Callaerts P, Norga K, Mackay TFC, Anholt RRH. 2008. Neurogenetic networks for startle-induced locomotion in *Drosophila melanogaster*. Proc Natl Acad Sci USA 105: 12393–12398.

Yamamoto A, Huang W, Anholt RRH, Mackay TFC. 2024. The genetic basis of variation in *Drosophila melanogaster* mating behavior. iScience, submitted.

